# Tailoring electrode surface charge to achieve discrimination and quantification of chemically similar small molecules with electrochemical aptamers

**DOI:** 10.1101/2022.06.25.497619

**Authors:** Vladimir Kesler, Kaiyu Fu, Yihang Chen, Chan Ho Park, Michael Eisenstein, Boris Murmann, H. Tom Soh

## Abstract

Electrochemical biosensors based on structure-switching aptamers offer many advantages because they can operate directly in complex samples and offer the potential to integrate with miniaturized electronics. Unfortunately, these biosensors often suffer from cross-reactivity problems when measuring a target in samples containing other chemically similar molecules, such as precursors or metabolites. While some progress has been made in selecting highly specific aptamers, the discovery of these reagents remains slow and costly. In this work, we demonstrate a novel strategy to distinguish molecules with miniscule difference in chemical composition (such as a single hydroxyl group) – with cross reactive aptamer probes - by tuning the charge state of the surface on which the aptamer probes are immobilized. As an exemplar, we show that our strategy can distinguish between DOX and many structurally similar analytes, including its primary metabolite doxorubicinol (DOXol). We then demonstrate the ability to accurately quantify mixtures of these two molecules based on their differential response to sensors with different surface-charge properties. We believe this methodology is general and can be extended to a broad range of applications.

## INTRODUCTION

In recent years, structure-switching electrochemical aptamer-based sensors^1–4^ have emerged as a powerful alternative to enzyme-based^5–8^ and antibody-based biosensing approaches.^8–10^ Such sensors typically entail tethering a redox label-modified aptamer to a gold electrode, wherein target binding alters the conformation of the aptamer and thereby generates a electrical signal change as an output. Such sensors can deliver low-cost, sensitive analyte detection while obviating the need for cumbersome sample preparation. There have been many innovations in selecting high-affinity aptamers using directed evolution,^11–14^ and also in improving the signaling of these aptamers by rationally designing switches^15,16^ and engineering electrode interfaces to tune transduction.^17–21^ However, one major challenge is that these sensors often suffer from poor specificity, such that the presence of structurally-similar interferents and metabolites can potentially make accurate molecular quantification of the target difficult or impossible. This is because the specificity of molecular detection is heavily dependent on the aptamer’s capacity to discriminate between chemically similar targets, and it can be exceedingly difficult to generate aptamers that exhibit such excellent selectivity while also achieving a robust target-induced conformational change.^14^

In this work, we report an alternative approach for improving target selectivity of electrochemical sensors based on structure switching aptamers. Our approach exploits the fact that electrochemical aptamer signaling depends on stochastic interactions between the redox reporter and the electric double layer (EDL) at the electrode surface.^22,23^ In recent work, we showed that the use of a nanoporous rather than a planar electrode improved the sensitivity of a sensor based on a doxorubicin (DOX) aptamer.^19^ This was because the highly-confined nanoporous structures extended the EDL, strengthening electrostatic effects near the electrode.^24,25^ We further hypothesized that altering the EDL composition could confer the benefit of introducing contrast between different targets. Specifically, we posited that tailoring the surface charge of the passivating self-assembled monolayer (SAM) would modulate the identity and concentration of counter-ions in the EDL. This would in turn change the sensor’s sensitivity to small chemical differences in the aptamer-target complex, thereby introducing target-specific signaling characteristics.

In this work, we apply these insights to demonstrate a sensor that can distinguish two molecules with miniscule differences in chemical composition, even when the aptamer cross-reacts between the two. As an exemplar, we show that our strategy can distinguish between DOX and many structurally similar analytes, including its primary metabolite doxorubicinol (DOXol) that only differs by a single hydroxyl group. We then demonstrate the ability to accurately quantify mixtures of these two molecules based on their differential response to sensors with different surface-charge properties. We designed multiple electrochemical aptamer sensors that produce a differential signal response by applying SAMs that incorporate charged end-groups. After confirming that tuning the EDL composition in this fashion affects the signaling of the DOX aptamer on planar electrodes, we then bolstered those effects by implementing the same sensor design on nanoporous electrodes. We demonstrated that altering the EDL composition endows excellent specificity, and subsequently engineered a sensor that can quantify both DOX and DOXol in the same sample by utilizing two electrodes with distinct SAMs and a mathematical model that describes the competitive binding of these molecules. This strategy should be broadly applicable to many aptamers and a wide range of targets, and could offer a valuable opportunity to expand the practical utility of electrochemical aptamer sensors for real-world molecular detection applications.

## RESULTS & DISCUSSION

### Modulating surface charge to tune aptamer performance

The readout from electrochemical aptamer-based sensors is often measured using square-wave voltammetry (SWV), a technique that measures the electron transfer rate due to stochastic collisions between an aptamer-conjugated methylene blue (MB) redox tag and electric fields in the EDL.^26,27^ The aptamer’s affinity and specificity determine the binding to a target analyte, but the subsequent signaling event depends on the difference in molecular dynamics between the unbound aptamer and the aptamer-target complex. This change in molecular dynamics and associated signal response is typically achieved by using aptamer switches that change structure upon binding.

Because electron transfer from MB to the electrode occurs at a fixed potential, surface charges determine the ionic composition of the EDL, which in turn affects the behavior of the aptamer-target complex as it approaches the electrode. For a planar electrode interface, the thickness of the EDL is described by the Debye length, which is approximately 1 nm under physiological conditions. However, on concave surfaces, the EDL extends farther into the solution because there is less space for ions to assemble at the interface. This is most easily understood by considering the “Debye volume,” which is the volume enclosed by the interface and an imaginary surface one Debye length normal to the interface.^24,25^ When the Debye volume-to-surface area ratio is low, the EDL extends further into the solution due to ionic crowding at the interface. In previous work, we showed that we could extend the EDL by employing nanoporous structured electrodes that constrain the Debye volume rather than conventional planar electrodes, thereby improving aptamer signaling and the sensitivity of the biosensor.^19^ Even with this improvement in signal strength and sensitivity, however, the specificity of the sensor remains dependent on the aptamer’s capacity to discriminate between target analytes and structurally-similar interferents.

Importantly, we hypothesized that local changes in ion concentrations can also affect the dynamics of signal transduction when the aptamer is bound to different targets. Altering the local ion concentrations at the interface translates into meaningful changes in aptamer signaling with a specific target,^28–30^ as do small changes in the charge or electrophoretic mobility of the complex due to a non-specific target.^31,32^ We inferred that interactions with ions in the EDL cause both phenomena, and we can leverage these effects to engineer specificity. We further posited that this happens because the signaling of an electrochemical aptamer is influenced by many secondary effects beyond aptamer binding. These effects include differences in the aptamer-target complex structure, interactions between the aptamer and the EDL, and interactions between the target and the EDL. Though it is hard to predict the impact of any particular change in the analyte, all these target-dependent effects influence the electron transfer rate. We therefore recognized a potential opportunity to engineer such specificity by tuning the properties of the EDL at the electrode interface. Traditionally, electrochemical aptamer-based sensor electrodes are passivated with SAMs featuring neutral end groups, but we hypothesized that changes in the identity and charge of those end-groups could affect electrochemical aptamer performance in a way that modulates sensor specificity.^28–30^

We therefore altered the composition of the passivating SAM on the electrode surface to modulate the surface charge at the interface. We first immobilized the aptamer onto the electrode via a gold-thiol linkage and then passivated the remaining electrode surface with one of two different alkyl-thiols: 6-mercapto-1-hexanol (C6-OH) (**Fig. 1A**) or 6-mercaptohexanoic acid (C6-COOH) (**Fig. 1B**). Alkyl-thiols terminated with end groups of low pKa value, such as C6-COOH, take on negative charges at physiological pH. In contrast, for C6-OH, where the end-group pKa approaches 7, the surface charge becomes neutral. By modifying the SAM in this fashion, we hypothesized that we could further build on the sensitivity gains achieved through the use of nanoporous structured electrodes to develop electrochemical aptamer sensors that simultaneously achieve excellent signal gain, linear range, and analyte specificity.

**Figure 1.**
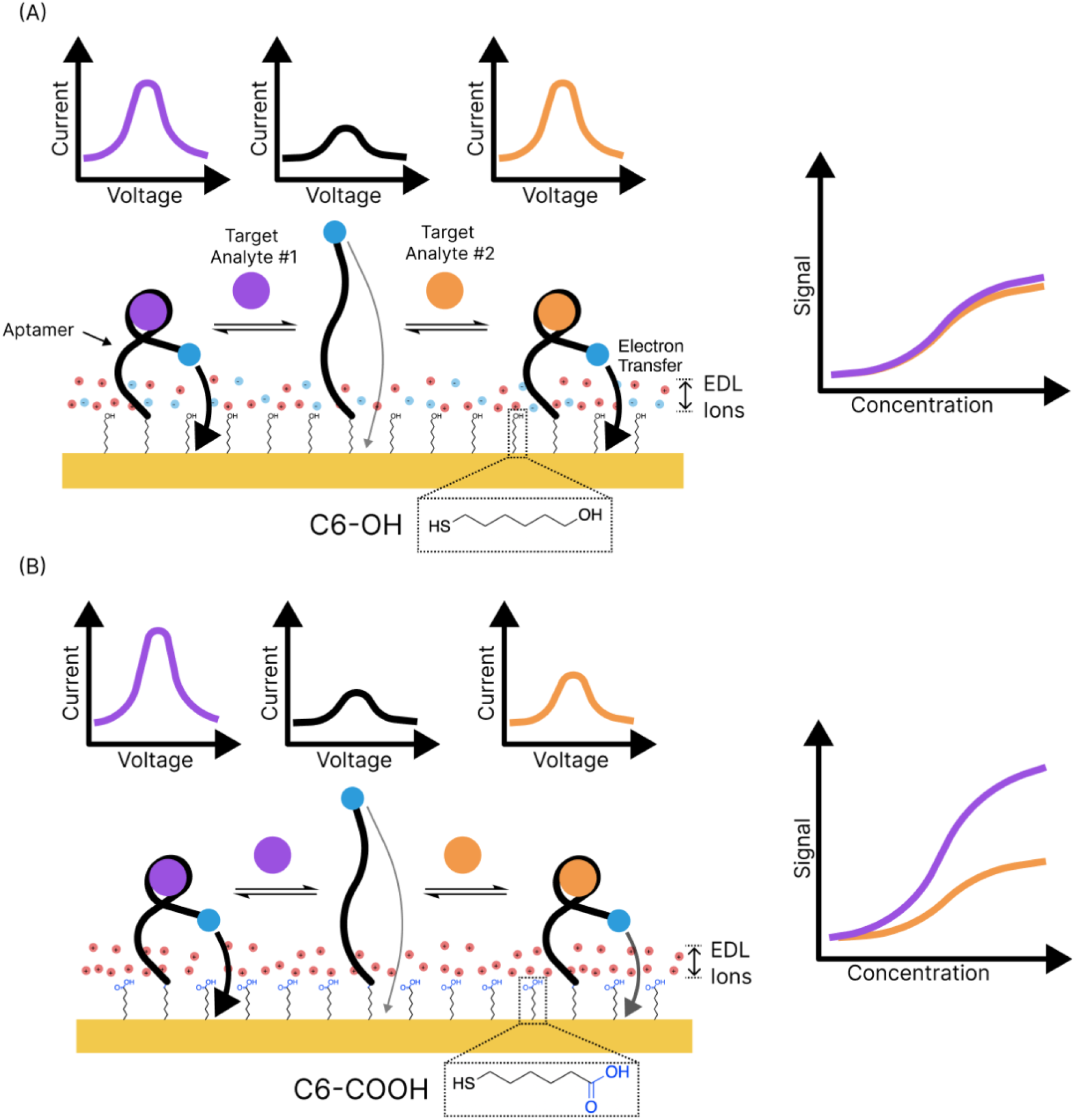
Schematic illustration of a dual electrochemical aptamer sensor for enhanced electrochemical biosensing performance. The aptamer is immobilized onto the electrode together with differently charged thiol molecules, which determine the ionic content of the EDL. (A) On the C6-OH passivated surface, the EDL assembles to balance neutrally charged end-groups (B) However, on the C6-COOH passivated surface, the negative charges from the carboxylic acid group alter the ionic composition of the EDL. This alters the signaling of the aptamer.

### Nanoporous electrodes enhance the effects of surface charge

We initially characterized the effects of SAM surface charge in the context of electrochemical aptamer sensors on planar electrodes. We generated binding curves by spiking-in DOX at a range of target concentrations and then converting the resulting SWV peak measurements to a signal gain metric. Signal gain is the baseline-subtracted ratio between the SWV peak generated in the presence of the target molecule to that produced in the absence of target; we used this value to quantify analyte concentrations in the sample.^3,19,33^ We found that the signal gain produced by 10 μM DOX (the maximum clinically relevant concentration) increased by ∼36% on the C6-OH SAM relative to the C6-COOH SAM—from 129% to 176% (**Fig. 2A**, upper plot). In order to quantitatively determine the change in sensitivity that resulted from this modification, we assessed the equilibrium dissociation constant (K_D_) of the sensor, which represents the midpoint of the linear range in the binding curve. After fitting the signal gain curve to a single-site binding isotherm and subsequently normalizing to the saturated value, we were able to measure a ∼34% decrease in K_D_ from 1.54 μM on the C6-OH surface to 1.01 μM on the C6-COOH surface (**Fig. 2A**, lower plot). This improvement in sensor affinity—and therefore in sensitivity—was entirely a product of the negative surface charge from C6-COOH, which confirmed that such changes in surface chemistry could sufficiently alter the EDL composition to affect electrochemical aptamer signaling. Notably, this enhanced sensitivity comes from improvements in both the sensor’s output dynamic range, which is the signal gain, and the sensor’s input dynamic range, which is the affinity.

**Figure 2.**
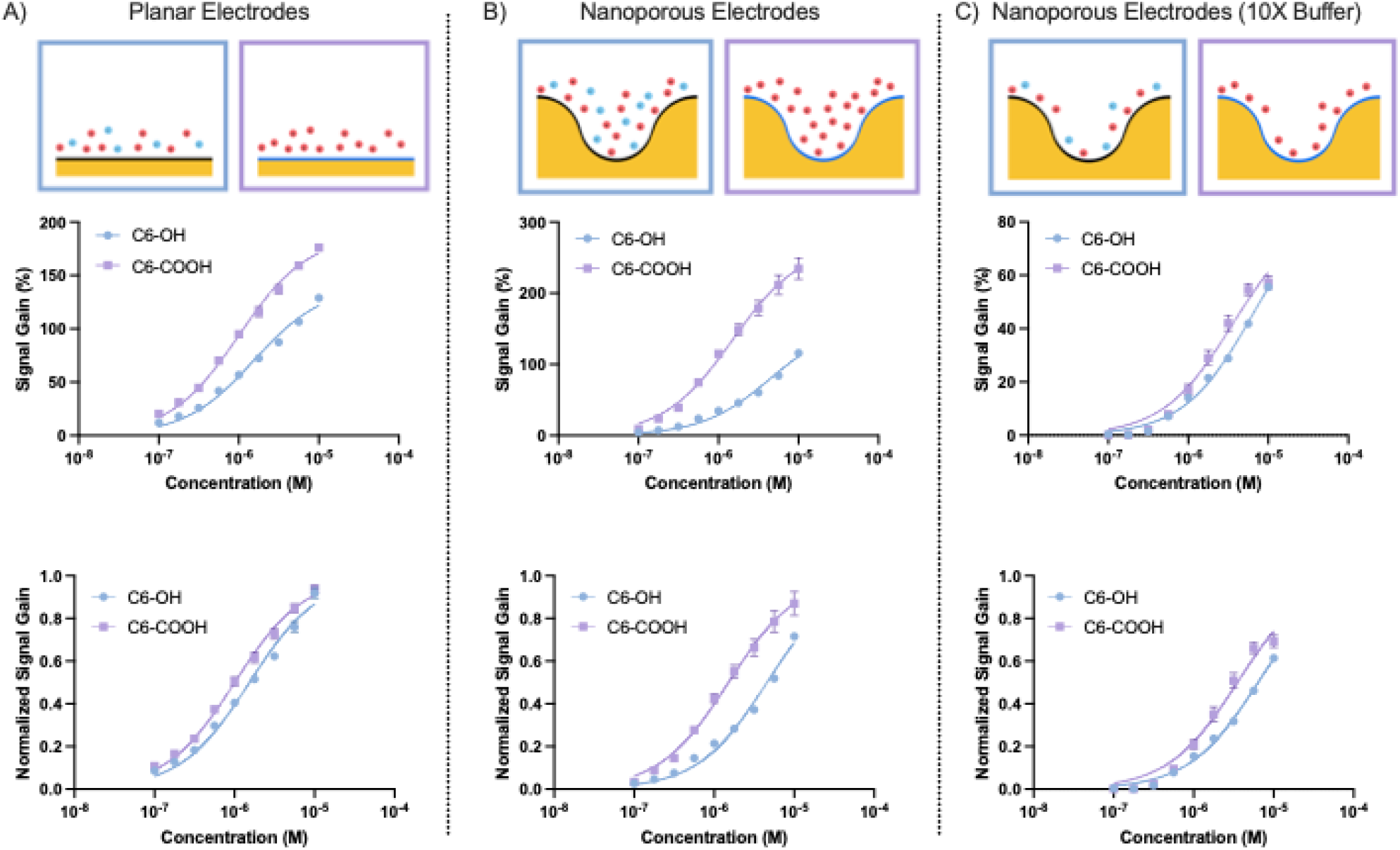
Effect of surface charge on DOX detection. Non-normalized (top) and normalized (bottom) DOX binding curves from sensors modified with C6-OH (blue) or C6-COOH (purple) on (**A**) planar electrodes, (**B**) nanoporous electrodes, or (**C**) nanoporous electrodes in 10X buffer. All plots are the average of three or four replicates; error bars represent the standard deviation (too small to be shown in some cases).

We hypothesized that we could further enhance the sensitivity of the sensor by using a nanoporous structured electrode that constrains the Debye volume of the electrode interface and thereby extends the EDL. And indeed, we observed a two-fold increase in signal gain from 116% on the C6-OH surface to 235% on the C6-COOH surface (**Fig. 2B**), in contrast to the ∼36% improvement seen on planar electrodes. After normalizing the binding curves, we saw that the K_D_ of the sensor shifted from 4.58 μM with C6-OH to 1.51 μM with C6-COOH—a three-fold improvement, versus the ∼34% shift seen on planar electrodes. These results confirmed our hypothesis, and demonstrated that the extended EDL produced by the concave morphology of the nanoporous electrode can further amplify the signal gains arising from the charged SAM. Based on these results, we solely employed nanoporous electrodes in subsequent experiments.

If this change in signaling is indeed the consequence of enhanced electrostatic interactions between the EDL and the MB tag on the aptamer switch, it should be possible to reverse these gains by establishing ionic conditions that contract the EDL. To test this, we repeated the above experiment using nanoporous sensors in highly concentrated (10X) buffer. Since the Debye length scales as 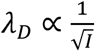, we predicted that a 10-fold increase in ionic strength (*I*) should essentially have the effect of shifting the EDL morphology on nanoporous electrodes to more closely resemble that seen with planar electrodes. As expected, the increased ionic strength minimized the differences between the two SAM conditions, with signal gains at 10 μM DOX of 56% and 57% for C6-OH and C6-COOH, respectively (**Fig. 3C**). After normalization, the reduction in K_D_ achieved with C6-COOH was just 45%—3.53 μM, versus 6.40 μM with C6-OH. Ultimately, the high ionic strength limits the reach of the EDL and erases the effects of the altered EDL composition. Overall, these results demonstrate that the signal enhancement we achieved above is a direct product of enhanced electrostatic interactions between the aptamer redox tag and the extended EDL.

**Figure 3.**
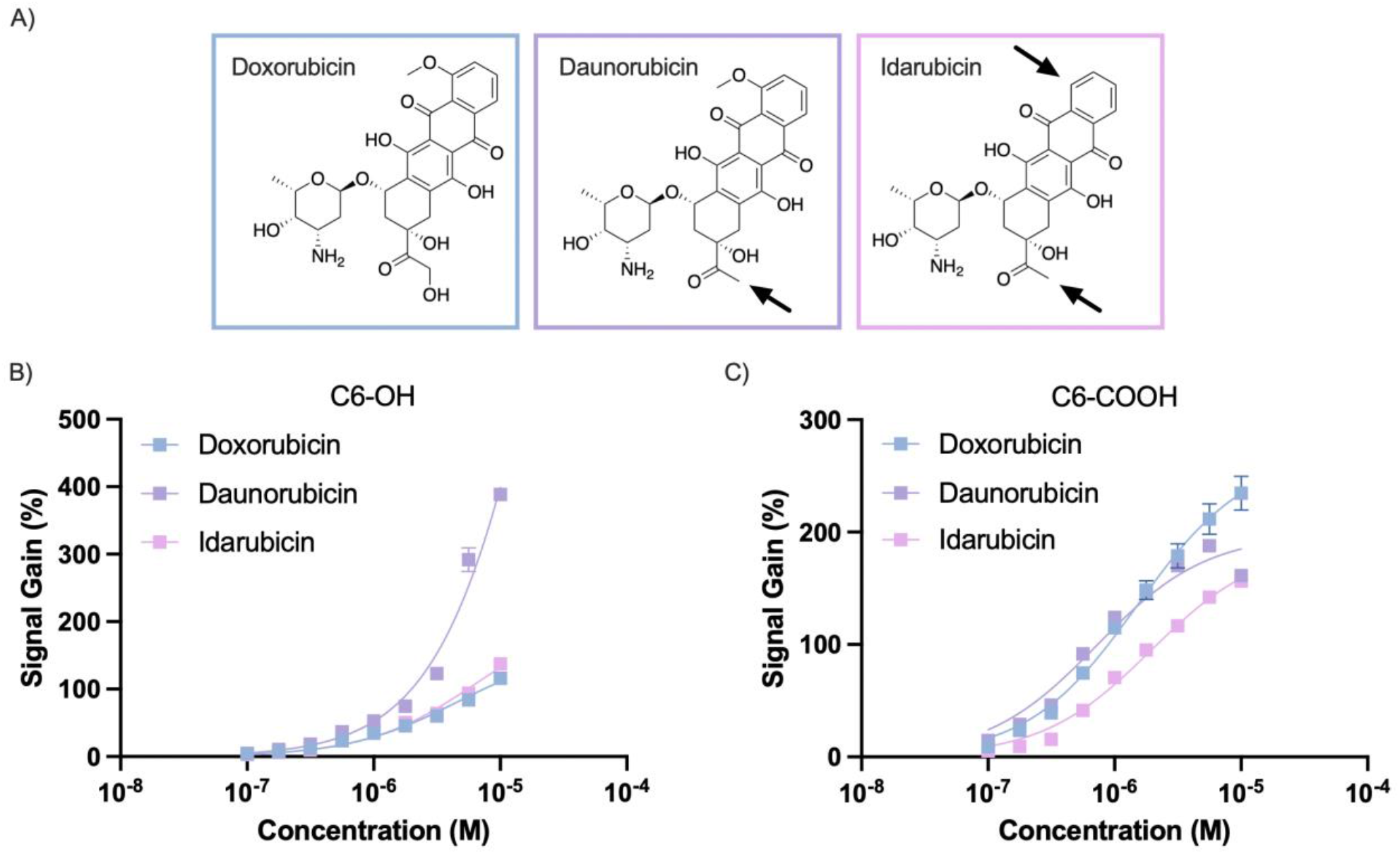
Different surface charge profiles affect the signal produced by different aptamer ligands. We tested (**A**) doxorubicin and two structurally similar analogs on sensors with (**B**) C6-OH and (**C**) C6-COOH surfaces. Plots are the average of three or four replicates; error bars represent the standard deviation (too small to be shown in some cases).

### Identification of chemotherapy drug analogs

We next examined the extent to which we could tune the sensor’s response to target versus non-target molecules by altering the EDL composition. As a test of sensor specificity, we employed daunorubicin and idarubicin—two structurally-similar analogues to DOX—as non-target interferents **(Fig. 3A)**. Daunorubicin is a dehydroxylated version of DOX, while idarubicin is missing an additional methoxy group, with both modifications occurring on the tetracycline ring. We chose these targets because our DOX aptamer cross-reacts with these two targets as hydrogen bonding between the solvent and the tetracyclic ring system at the core of these molecules has been implicated in stabilization of the aptamer-target complex.^34^ Because idarubicin has a further modification beyond daunorubicin, we expected that any changes in signaling relative to DOX would be further enhanced for idarubicin. To test this theory, we again prepared sensors using C6-OH and C6-COOH SAMs, and generated binding curves against these new targets.

When we tested the sensors, we confirmed our initial hypothesis that different analytes will produce distinct changes in signaling depending on the surface modification, although the nature of these changes was not predictable *a priori*. On the C6-OH surface, the sensor responded identically to DOX and idarubicin, while daunorubicin produced an amplified response (**Fig. 3B**). In contrast, DOX and daunorubicin produced similar responses on the C6-COOH surface in the clinical concentration range, with a weaker response to idarubicin (**Fig. 3C**). These results are compelling because these chemotherapy drug analogs have highly similar chemical structures, and thus demonstrate that the mechanism of transduction of electrochemical aptamer sensors enables the discrimination of target analytes. Further, these results highlight the complexity of the mechanism of signaling for electrochemical aptamer sensors: rather than amplifying the signal alterations observed with daunorubicin relative to DOX, idarubicin’s additional modification reversed the enhanced signaling seen with daunorubicin on the C6-OH surface and resulted in suppressed signaling relative to the other two analytes on the C6-COOH surface.

Our data also show that differences in surface charge affect both the signal transduction and the binding affinity of the aptamer to the different targets. We can illustrate the influence on signal transduction by looking at the *B*_*max*_ on our fitted single-site binding isotherms, which describes the saturated signal gain value of the binding curve. For example, on the C6-OH surface, daunorubicin has by far the highest *B*_*max*_, at 1,554%, versus 216.7% for idarubicin and 162% for DOX. Meanwhile, the K_D_ of the sensor is an order of magnitude higher for daunorubicin (28.9 μM) compared to 4.58 μM for DOX and 6.42 μM for idarubicin (**Fig. S1A**). Thus, the differences in the clinical concentration range are dominated by the influence of signal transduction, despite the dramatic increase in K_D_. Signal transduction and binding affinity both play a role on the C6-COOH surface as well. While *B*_*max*_ decreases from 269.7% for DOX to 198.3% for daunorubicin and 189.9% for idarubicin, K_D_ decreased from 1.51 μM for DOX to .72 μM for daunorubicin but increased to 1.95 μM for idarubicin (**Fig. S1B**). In this scenario, both daunorubicin and idarubicin produce altered signal transduction compared to DOX, but the change in affinity for daunorubicin counters this effect and makes the signals appear similar in the measured range. Given this differential response, it is clear that by obtaining access to measurements from both surfaces, it should become straightforward to discriminate DOX from its analogues.

### A differential sensor pair for metabolite sensing

Having established this ability to discriminate highly similar analytes, we extended this strategy to develop a dual sensor for DOX and its primary metabolite DOXol. Accurate dosing of DOX can minimize the severe side effects of this chemotherapy drug,^35^ but individuals metabolize DOX at different rates and correcting for the presence of DOXol has proven challenging because these two molecules can currently only be discriminated with low-throughput, high-cost methods like mass spectroscopy and HPLC.^36,37^ DOXol is particularly challenging to discriminate from DOX, as it differs from its parent molecule by only one −OH group (**Fig. 4A**). Both targets produced a similar signal gain response on the C6-OH surface (**Fig. 4B**), but could readily be discriminated on the C6-COOH surface (**Fig. 4C**). With the latter sensor, the signal gain at 10 μM was 100% for DOXol versus 235% for DOX, with K_D_ values of 2.56 μM and 1.51 μM, respectively (**Fig. S2**).

**Figure 4.**
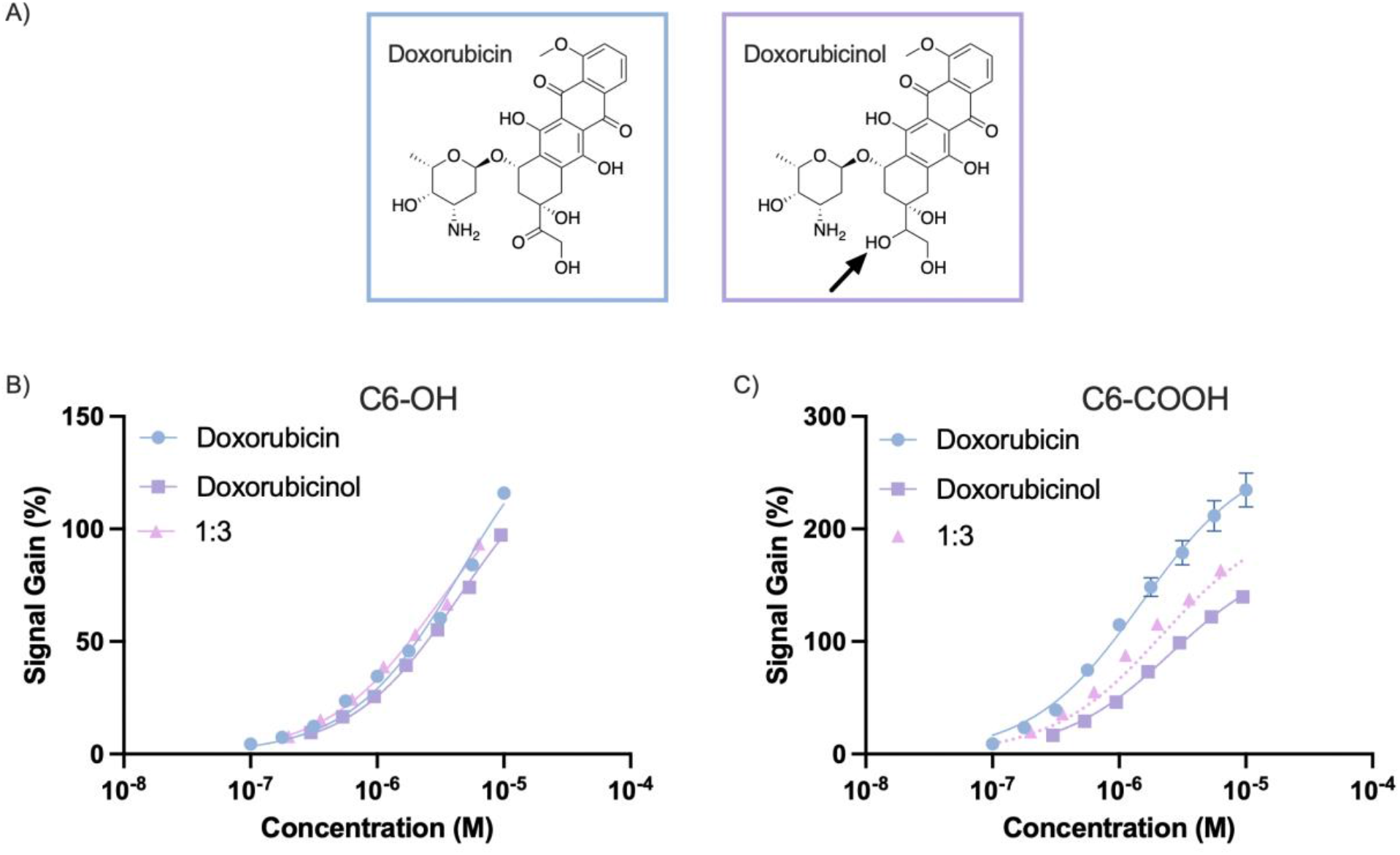
Differential responses of our electrochemical sensor to DOX and DOXol with different SAMs. (A) The two structures differ by only one hydroxyl group, indicated by the arrow. (B, C) We tested our sensors against DOX (blue), DOXol (purple) and a 1:3 mixture of DOX and DOXol (pink) on sensors passivated with (B) C6-OH and (C) C6-COOH. Solid lines are fitted binding isotherms, dotted lines are predicted curves from a competitive binding model. Error bars (some too small to display) represent the standard deviation of n = 3 (DOX) or n = 4 (DOXol, 1:3 DOX:DOXol) replicates.

Importantly, these differences in signaling can be used to quantify the concentrations of both DOX and DOXol when present as mixtures with unknown ratios in a given sample. This is done by using the data from the C6-OH surface to “anchor” the total concentration of analyte, and using the data from the C6-COOH surface to identify the proportion of the analyte that is DOX or DOXol. To achieve this, we leveraged the well-understood thermodynamic relationships that govern competitive binding of DOX and DOXol to the DOX aptamer. Importantly, these relationships depend only on the values for *K*_*D*_ and *B*_*max*_, and we were able to extract these values by fitting single-site isotherms to the binding curves for each target on each surface. Next, we derived the binding fractions, denoted Θ, of DOX and DOXol across concentrations from first principles^38^:

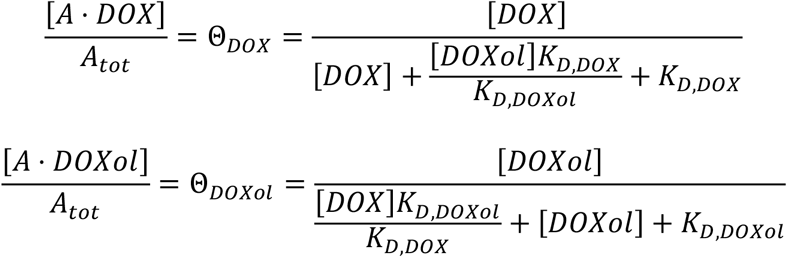

We then converted these binding fractions to signals in a target-dependent manner:

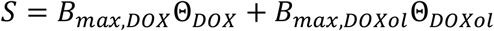

By taking measurements using the sensor pairs, we can establish a system of two non-linear equations with two unknowns: the analyte concentrations. We can thereby predict the signals produced when various arbitrary mixtures of DOX and DOXol are present in a sample. To illustrate this capability, we tested a 1:3 DOX:DOXol cocktail, and overlaid the competitive binding model’s prediction for the signals (**Fig. 4C**).

This model’s usefulness comes not from predicting unknown signals from known concentrations, but rather from quantifying unknown molecular concentrations from measured signals. In order to demonstrate that we can quantify molecular concentrations using our model, we generated calibration heat maps. By using the competitive binding relationships to solve for DOX and DOXol concentrations across the full range of signal gain from each sensor, we obtained a three-dimensional calibration curve for each target using a non-linear numerical equation solver (**Fig. 5A, B**). On the C6-OH surface, this range was from 1–150%, and on the C6-COOH surface, it spanned from 1–250%. Signal gains that cannot be achieved with real-world DOX and DOXol concentrations were omitted from the heat map. We then demonstrated the robustness of our predictive model using the signal gain measurements from the 1:3 cocktail of DOX and DOXol from both sensors. We put this data through the calibration map and plotted it against the true concentrations in order to see how well the predictions correlated with our experimental results (**Fig. 5C)**. Rather than reporting mean values and deviations, we used all the combinations of our four replicates on each surface, yielding 16 binding curves per target. We determined strong correlation (r > .984) for all the data across both targets relative to the actual concentrations of the drug and its metabolite. Collectively, these data demonstrate that a competitive binding model derived from the binding curves for individual targets can accurately quantify arbitrary combinations of DOX and DOXol in the same sample. Furthermore, these results show that it is possible to tailor electrode design in order to achieve excellent discrimination of closely related analytes from an electrochemical aptamer sensor, even when the aptamer itself lacks that level of innate selectivity.

**Figure 5.**
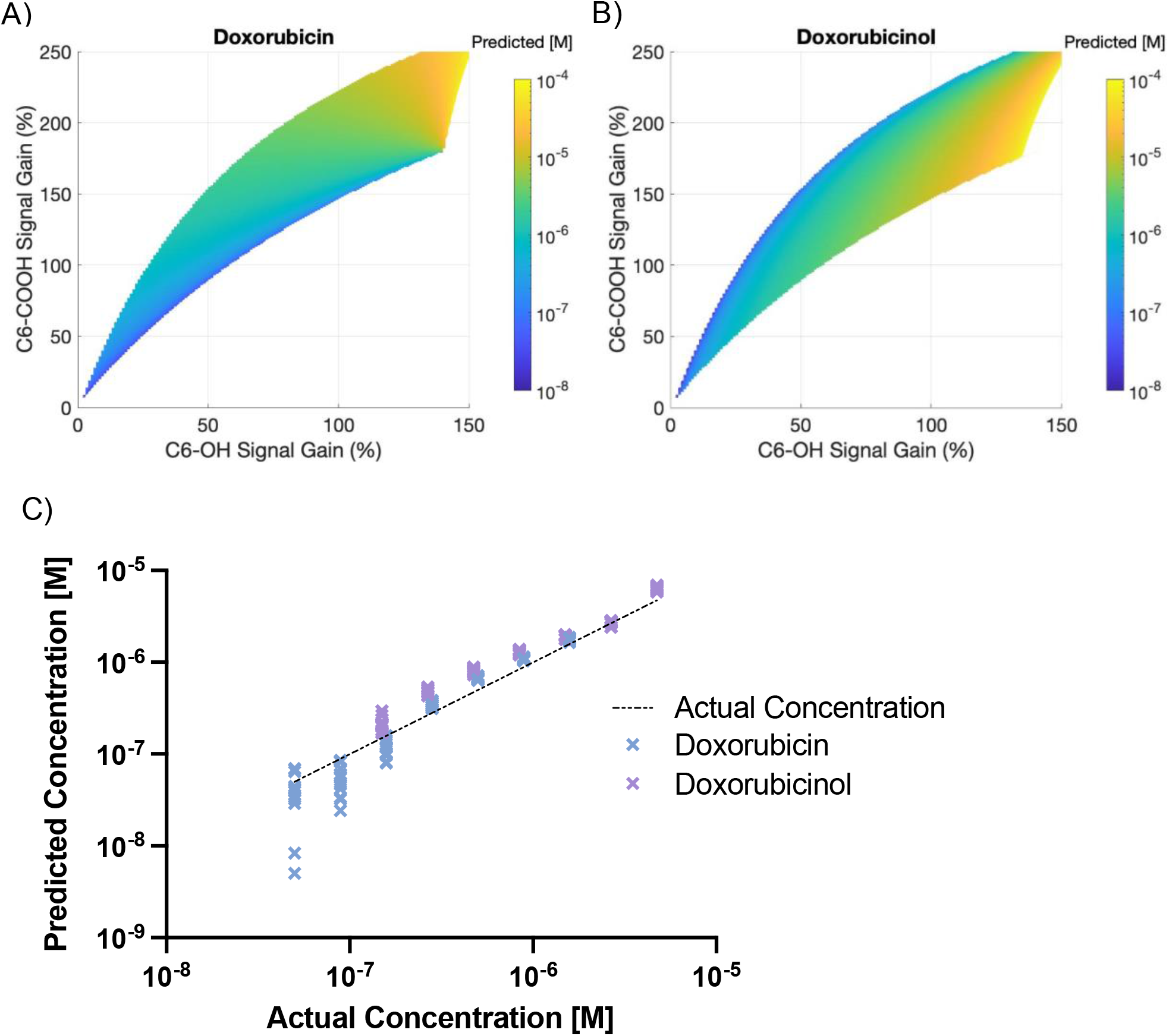
Achieving robust quantitation of DOX:DOXol mixtures based on a predictive model derived from individual analyte measurements. We generated calibration heat maps for the signal output of both sensors in response to (A) DOX and (B) DOXol. (C) We then ran data from a 1:3 mixture of DOX (blue) and DOXol (purple) through this map, and plotted the results against their true concentrations (Data represents all combinations of four replicates on each surface, yielding 16 binding curves).

## CONCLUSION

Electrochemical aptamer-based sensors can achieve excellent sensitivity, even in complex samples, but their specificity is generally constrained by the selectivity of the aptamer that is being employed. In this work, we demonstrate an alternative strategy for achieving such discriminatory power, in which we modify the surface charge of electrodes to tailor the EDL—and thus the electrochemical response produced in response to distinct analyte-binding events. We have shown that by incorporating charged end-groups in the passivating SAM applied to the sensor surface, we can change the ionic composition of the double layer and improve the signal gain of the electrochemical DOX aptamer-based sensor. These effects can be further enhanced by using nanoporous structured electrodes rather than planar electrodes, thereby constraining the Debye volume and extending the thickness of the EDL. We show that these differential surface-charge effects can be used to distinguish chemically similar analytes that cross-react with the same aptamer, and developed a predictive model that can be used to accurately quantify undefined mixtures of two closely-related analytes—in this case, DOX and DOXol. This represents an important opportunity for the development of electrochemical sensors with excellent discriminatory power, as it obviates the need to laboriously isolate, engineer, and optimize aptamers that produce the desired target-specific response profile.

While we have demonstrated that the EDL can be engineered to alter the signal transduction and enable specific quantification of multiple analytes, there remain many unanswered questions regarding the mechanisms underpinning this phenomenon because there are many effects that can impact the out signal. This includes electrostatic interactions between the EDL and the aptamer, the EDL and the analyte, as well as changes in the structure of the aptamer-target complex due to chemical differences in the analytes. Although it is beyond the scope of current work, further studies are needed to understand the relative contributions that each of these effects have on the output signal. Once these mechanistic details have ben elucidated, we foresee opportunities in developing highly multiplexed biosensor systems for wide range of applications including monitoring of metabolic pathways, synthesis reactions, and testing the purity of small-molecules, without the need for highly specific aptamers.

## MATERIALS AND METHODS

### Reagents and Materials

The DOX aptamer was adapted from previous work^3,33^ and purchased from Integrated DNA Technologies: 5’-S-S-ACCATCTGTGTAAGGGGTAAGGGGTGGT-NH_2_-3’. Methylene blue (MB) was attached to the 3’ terminus by conjugating the aptamer with a MB-NHS ester (Biosearch Technologies), followed by ethanol precipitation. Reaction yield was quantified via UV-Vis spectrophotometry (Nanodrop 2000, Thermo Fisher Scientific). 6-mercapto-1-hexanol (C6-OH), 6-mercaptohexanoic acid (C6-COOH), tris(2-carboxyethyl)phosphine (TCEP), DOX, daunorubicin and idarubicin were obtained from Sigma-Aldrich. Doxorubicinol was obtained from Santa Cruz Biotechnologies. 1X SSC buffer solutions were prepared by diluting a stock solution (20X, pH 7.4, Thermo Fisher Scientific) in MilliQ deionized (DI) water. For the reference electrode, silver wire was obtained from A&M Systems, and coated with AgCl by incubating in bleach (Clorox) for 15 minutes, followed by gentle cleaning in 70% ethanol. Platinum counter electrodes were purchased from CH Instruments.

### Device Fabrication

Electrode arrays were patterned on Pyrex wafers using a standard lift-off process with Shipley 3612 resist. For nanoporous electrodes, Ti (10 nm) and Au (50 nm) metal films were deposited via a sputtering process to form a base layer, followed by co-sputtering of a Ag:Au alloy (2:1 ratio, 300 nm). Next, the Ag was wet etched out of the alloy in nitric acid (70% v/v) at room temperature for 16 hours. Following lift-off, the electrodes were encapsulated in SU-8 3005 (Kayuku) to make 1 mm x 1 mm active areas for the sensors. Electrodes were then characterized using SEM/EDS **(Figure S3)**. For planar electrodes, Ti (10 nm) and Au (90 nm) films were deposited via e-beam evaporation (AJA). No wet etching step was necessary before encapsulation for the planar electrodes. Polydimethylsiloxane (PDMS) wells for the solutions were placed around the electrodes. Immediately prior to functionalization, the electrodes were cleaned electrochemically by cyclic voltammetry in sulfuric acid (10 scans in 200 mM, 10 scans in 50 mM) followed by thorough washing in DI water.

Electrode devices were then functionalized with the electrochemical aptamer and then a passivating SAM. First, 100 μM aptamer (in DI) was incubated with TCEP (28.7 mg/mL in DI) at a 1:2 ratio for 40 minutes in the dark. The aptamer solution was then diluted to 1 μM in 1X SSC buffer. Immediately afterwards, electrodes were dried and immersed in the aptamer solution for 1 hour. For passivation, 10 mM C6-OH and C6-COOH solutions were prepared in 1X SSC buffer. The electrodes were washed thoroughly in 1X SSC buffer and incubated for 2 hours in C6-X solution. Finally, electrodes were washed again in 1X SSC, and stored in 1X SSC overnight at room temperature in the dark before measurements.

### Electrochemical Aptamer Sensor Characterization

All measurements were performed in 1X SSC using an emStat3blue potentiostat. SWV was carried out at 200 Hz with the potential range from −0.5 to −0.1 V with an amplitude of 20 mV and a step size of 2 mV. Binding curves were generated by titrating concentrated stock solutions of target analytes (in SSC buffer) into the PDMS well, pipette mixing thoroughly, and waiting 10 minutes for equilibration. SWV peak fitting was done with a custom MATLAB script; signal gain was calculated by referring SWV peaks to the target-free buffer condition as follows:

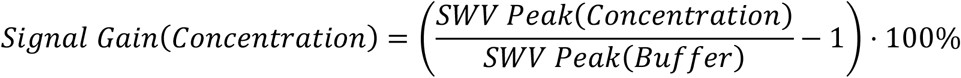

*K*_*D*_ and *B*_*max*_ were subsequently extracted by fitting the signal gain data to a single-site specific binding isotherm in Graphpad Prism 9.

## Supporting information

Fu_Seo_Kesler_SI

## ACKNOWLEDGEMENTS

This work was supported by the Chan-Zuckerberg Biohub, the Helmsley Trust, W.L Gore and Associates, and the National Institutes of Health (NIH, OT2OD025342). Device fabrication was performed at the Stanford Nanofabrication Facility (SNF) and SEM/EDS characterization was performed at the Stanford Nano Shared Facilities (SNSF), which are supported by the National Science Foundation under award ECCS-2026822. The authors thank Liwei Zheng, Alexander Yoshikawa and Brandon Wilson for helpful discussions.

## AUTHOR CONTRIBUTIONS

V.K., K.F. and H.T.S. designed the research. V.K. and Y.C. fabricated and characterized the devices. V.K. and K.F. designed the experiments. V.K. and K.F. executed the experiments, collected the data, and analyzed the data. V.K., K.F. and H.T.S. wrote the paper. All authors discussed the data and edited the paper.

