## Supplementary material for "Tailoring electrode surface charge to achieve discrimination and quantification of chemically similar small molecules with electrochemical aptamers": Fu_Seo_Kesler_SI

### SUPPLEMENTARY INFORMATION

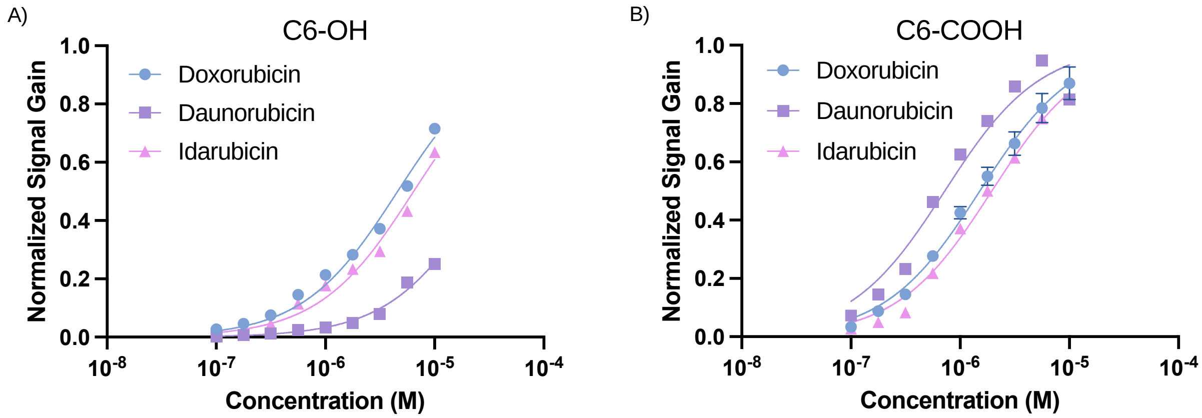


**Figure S1**. Normalized binding curves of DOX and drug analogs daunorubicin and idarubicin on (A) the C6-OH surface and (B) the C6-COOH surface. Plots are the average of three or four replicates; error bars represent the standard deviation (too small to be shown in some cases).

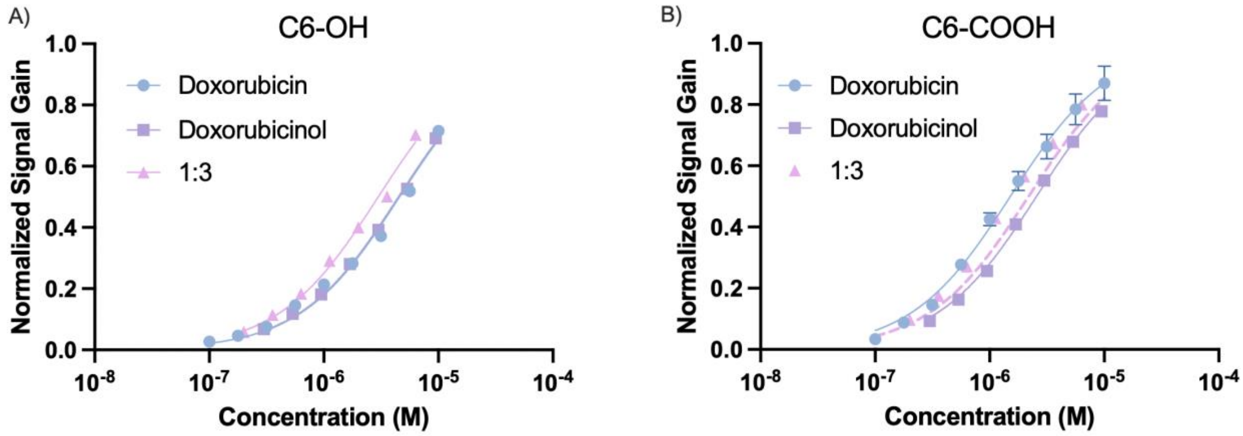


**Figure S2**. Normalized binding curves of DOX, DOXol, and a 1:3 cocktail of the drug and its metabolite on (A) the C6-OH surface and (B) the C6-COOH surface. Plots are the average of three or four replicates; error bars represent the standard deviation (too small to be shown in some cases).

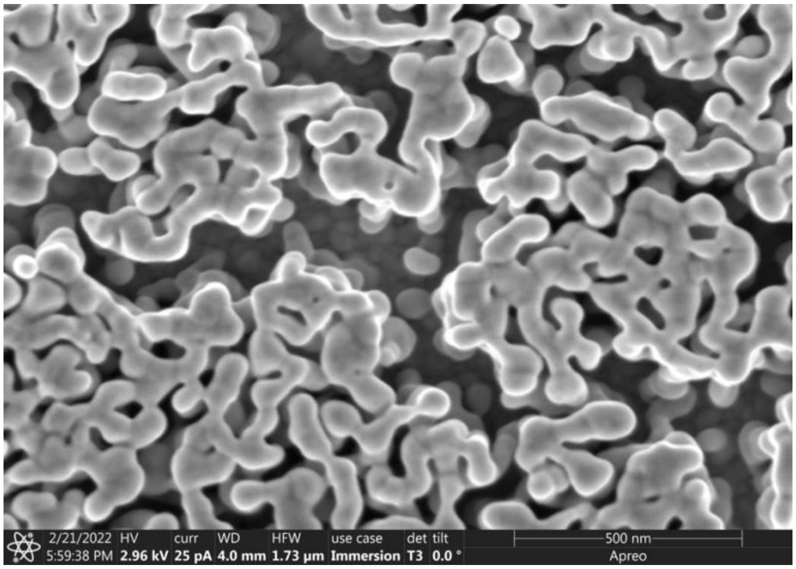


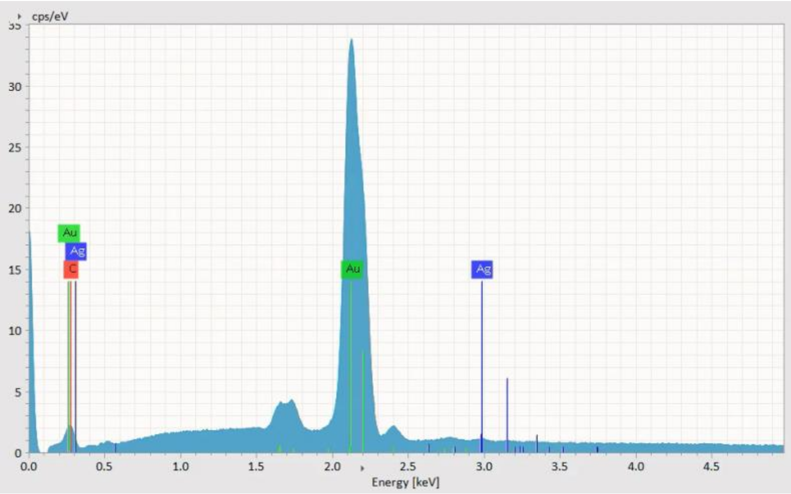


**Figure S3**. SEM (top) and EDS (bottom) characterization of representative nanoporous gold electrodes. EDS analysis shows minimal Ag content remaining after selective wet etching, as well as Ti peaks from the adhesive metal layer and Si, O, and Na peaks from the glass substrate.
